# Accucopy: Accurate and Fast Inference of Allele-specific Copy Number Alterations from Low-coverage Low-purity Tumor Sequencing Data

**DOI:** 10.1101/2020.01.02.892364

**Authors:** Xinping Fan, Guanghao Luo, Yu S. Huang

## Abstract

**Background:** Copy number alterations (CNAs), due to their large impact on the genome, have been an important contributing factor to oncogenesis and metastasis. Detecting genomic alterations from the shallow-sequencing data of a low-purity tumor sample remains a challenging task.

**Results:** We introduce Accucopy, a method to infer total copy numbers (TCNs) and allele-specific copy numbers (ASCNs) from challenging low-purity and low-coverage tumor samples. Accucopy adopts many robust statistical techniques such as kernel smoothing of coverage differentiation information to discern signals from noise and combines ideas from time-series analysis and the signal-processing field to derive a range of estimates for the period in a histogram of coverage differentiation information. Statistical learning models such as the tiered Gaussian mixture model, the Expectation-Maximization (EM) algorithm, and Sparse Bayesian Learning (SBL) were customized and built into the model. Accucopy is implemented in C++/Rust, packaged in a docker image, and supports non-human samples, more at http://www.yfish.org/software/.

**Conclusions:** We describe Accucopy, a method that can predict both TCNs and ASCNs from low-coverage low-purity tumor sequencing data. Through comparative analyses in both simulated and real-sequencing samples, we demonstrate that Accucopy is more accurate than Sclust, ABSOLUTE, and Sequenza.

## Background

Genomic alterations discovered in large-scale cancer genomic projects [1, 2], have therapeutic implications in being an important source of drug development [3, 4]. Copy number alterations (CNAs), due to its large impact on the genome, have been an important contributing factor to oncogenesis and metastasis [5]. Increasingly, clinical tumors of varying purity, which is the fraction of tumor cells in a sample, are being characterized by bulk whole-genome sequencing (WGS). Different approaches have been applied to infer CNAs from these tumor sequencing data [6–10] but few automated and efficient methods exist to infer CNAs by shallow-sequencing low-purity tumor samples.

CNA-detection methods usually leverage two types of differentiating information: (1) the total sequencing coverage differentiation between the tumor and its matching normal sample, and (2) the allelic sequencing coverage differentiation between the two distinct alleles of heterozygous germline single-nucleotide variants (HGSNVs). The first type of differentiation is a predictor for total copy number (TCN). If the TCN estimate for a genomic region is not two, it indicates the presence of somatic copy number alterations (SCNAs). The second type of differentiation is a predictor for allele-specific copy number (ASCN). The ASCN estimates can help to discern more SCNAs, such as loss of heterozygosity (LOH). Based on how these two types of differentiation are utilized, existing computational methods can be broadly grouped into three categories. Category one utilizes total coverage differentiation only [11, 12]. Category two utilizes allelic coverage differentiation only [13, 14]. Category three utilizes both information [7, 9, 10, 15–18]. Some computational methods have ventured into inferring the evolutionary phylogeny [15, 17, 19, 20] from a single sample or multiple samples of the same tumor, Table 1. Although inferring the evolutionary phylogeny is an exciting subject, our method, Accucopy, aims to address another problem prevalent in clinical settings.

**TABLE 1.** Comparison of copy-number and subclonal architecture inference methods.

|  | Call<br>TCN | Call<br>ASCN | Subclonal<br>composition | Multi-<br>samples | Publish<br>year | Year of<br>last | Low<br>coverage | Low<br>Purity |
| --- | --- | --- | --- | --- | --- | --- | --- | --- |

|  |  |  |  |  |  | release |  |  |
| --- | --- | --- | --- | --- | --- | --- | --- | --- |
| ABSOLUTE[7] | Y | Y | N | N | 2012 | 2012 | N | N |
| **Accucopy** | Y | Y | N | N | * | 2020 | Y | Y |
| Canopy[15] | Y | Y | Y | Y | 2016 | 2017 | N | N |
| CITUP[19] | N | N | Y | Y | 2015 | 2016 | N | N |
| cloneHD[17] | Y | Y | Y | Y | 2014 | 2014 | N | N |
| CNAnorm[11] | Y | N | N | N | 2011 | 2020 | Y | N |
| PhyloWGS[20] | N | N | Y | Y | 2015 | 2019 | N | N |
| Sclust[9] | Y | Y | Y | N | 2018 | 2019 | N | N |
| Sequenza[10] | Y | Y | N | N | 2015 | 2019 | N | N |
| THetA[12] | Y | N | Y | N | 2015 | 2015 | N | N |
| TITAN[18] | Y | N | Y | N | 2014 | 2014 | N | Y |

N: no. Y: yes. \*: unpublished.

Collaboration with oncology clinicians provided us one of the impetuses to develop Accucopy: the large amount of discarded low-quality (i.e. low-tumor-purity) clinical samples because of the absence of a good computational method to extract clinically meaningful information (i.e. TCNs and ASCNs) from these low-purity samples. Clinicians are also reluctant to invest in high-coverage tumor sequencing outright, which is required to robustly infer the evolutionary phylogeny of a tumor in order to guide the next round of treatment. Low-coverage sequencing is an efficient alternative to rescue those low-quality samples. We observed the periodic patterns in the histogram of the total coverage differentiation information and thought they may offer a route to develop a better computational method. Previously, we have developed Accurity [21], a method that focuses on inferring tumor purity by using these differentiation information. In this manuscript, we describe Accucopy, a method to infer TCNs and ASCNs from challenging low-purity and low-coverage tumor samples. Accucopy adopts many robust statistical techniques such as kernel smoothing of coverage differentiation information to discern signals from noise and combines ideas from time-series analysis and the signal-processing field to derive a range of estimates for the period in a histogram of coverage differentiation information. Statistical learning models such as the tiered Gaussian mixture model, the Expectation-Maximization (EM) algorithm, and Sparse Bayesian Learning (SBL) were customized and built into the model. We also invested considerable efforts in making the software easy to use. Packaged in a docker image, Accucopy is highly automated and compatible to virtually all operating systems, even supporting non-human samples (online communication with other users). The result is an easy-to-use software that works much better than its peers do in low-coverage and low-purity tumor samples.

Cancer computational methodology is a fast-moving subject and we chose the latest or still-widely-used methods for comparative analyses. ABSOLUTE [7], is one of the most widely-used tumor copy number inference software despite its early publication. Its strength is that it can work with both array-based copy number data and sequencing data and can also use segmented copy number data derived from whole genome or exome sequencing. However, its performance in low-coverage and low-purity samples is questionable and it contains many manual steps. Sequenza [10], another widely-used recent method, employs a probabilistic model built upon the average depth ratio (tumor versus normal) and B allele frequency for each segment. One of its weaknesses is that copy number two is much preferred over other values via a prior probability function, which bodes ill for tumor genomes that underwent significant disruptions, i.e. whole-genome disruptions (WGDs). Sclust [9] is a fully nonparametric mutational clustering method that infers TCNs and ASCNs with low computational burden by using smoothing splines. Although its results were impressive, all samples analyzed in the publication are at least 30X coverage. Through comparative analyses in both simulated and real-sequencing samples, we demonstrate that Accucopy is more accurate than these methods, esp. in low-coverage and low-purity samples.

Next, we first describe the data and the Accucopy model. Then, we evaluate Accucopy on numerous simulated and real-sequencing samples and compare Accucopy with Sclust [9], Sequenza [10] and ABSOLUTE [7]. We end the paper with discussions on the strengths and weaknesses of Accucopy.

## Results

### Evaluation of Accucopy on the simulated data

We evaluated the FullC and CallF of Accucopy using simulated tumor and normal data under three coverage settings: 2X, 5X, and 10X, and nine different purity settings, 0.1-0.9, (Fig. 1). For the TCN inference, Accucopy achieved high FullC and CallF, mostly >0.95, regardless of the tumor purity level, even if the coverage is only 2X (Fig. 1A, 1C and 1E). In the low-purity (0.1-0.4) cases of the 2X coverage, the TCN FullC deteriorates only slightly to about 0.9. For the ASCN inference, Accucopy achieved robust FullC and CallF, >0.8, when the sample purity is equal to or above 0.2 (Fig. 1B and 1D) for the 5X and 10X coverage. In the low coverage settings (2X), Accucopy requires the purity of sample to be at least 0.6 to achieve good performance in ASCN inference, but the total copy number (TCN) estimates are still >90% correct (Fig. 1E). We think the extremely low tumor content (<0.4), less than 1.2X (=0.6*2X) coverage on average for the tumor cells, renders the ASCN inference quite challenging.

**Fig. 1.**
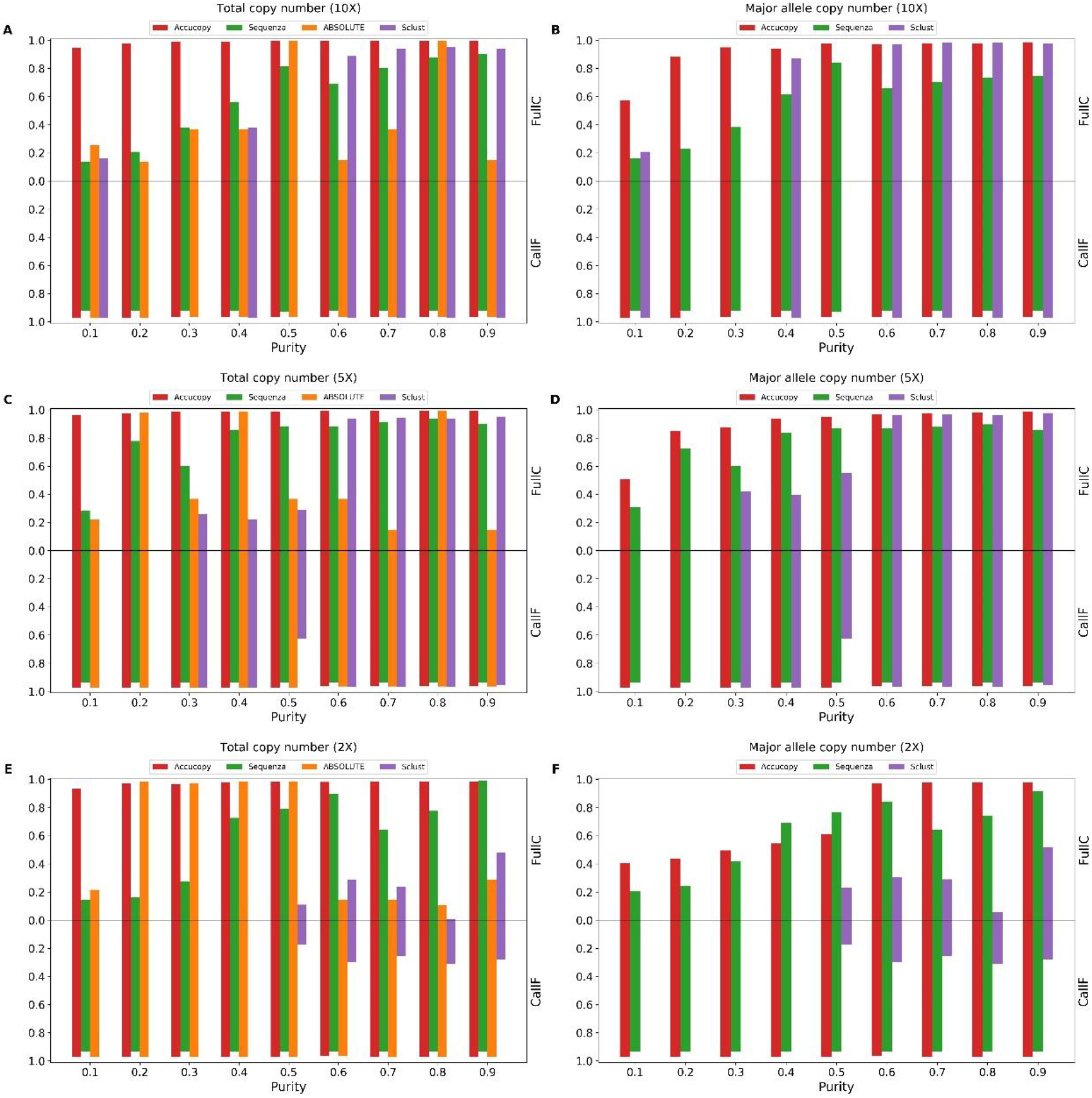
Evaluation of Accucopy and Sclust on simulation data. FullC and CallF of total copy number on low-coverage 10X (**A**), low-coverage 5X (**C**) and low-coverage 2X (**E**). FullC and CallF of major allele copy number on low-coverage 10X (**B**), low-coverage 5X (**D**) and low-coverage 2X (**F**). The blank space in the figure indicates Sclust failed on this sample. The red, green, orange and purple bar represent Accucopy, Sequenza, ABSOLUTE and Sclust respectively.

We compared Accucopy with Sclust, Sequenza, and ABSOLUTE. All three methods can infer the tumor purity, SCNAs, and ASCNs. Sclust performs well in high-coverage (>=10X) and high-purity settings (purity >=0.6) (Fig. 1A and 1B) and performs reasonably well in medium coverage (5X) and high purity settings (purity ≥ 0.6), but performs poorly in low-purity (<=0.5) or low-coverage (2X) settings. Sequenza performs similarly to Sclust in 5X and 10X but outperforms Sclust in 2X and low-purity conditions. Sequenza has the strange phenomenon that it performs better on 5X samples than 10X samples. We found that Sequenza over-segments the genome in 10X samples and calls many of these small segments with the wrong copy number and thus has lower performance in 10X than 5X. The case of ABSOLUTE is curious. It achieves good TCN performance on par with Accucopy in some conditions but performs quite poorly in other conditions. We found that its TCN performance is dependent on its ability to estimate the tumor purity correctly, (Table 2). Across the coverage and purity level, Accucopy is the top performer or a very close second.

**Table 2.**
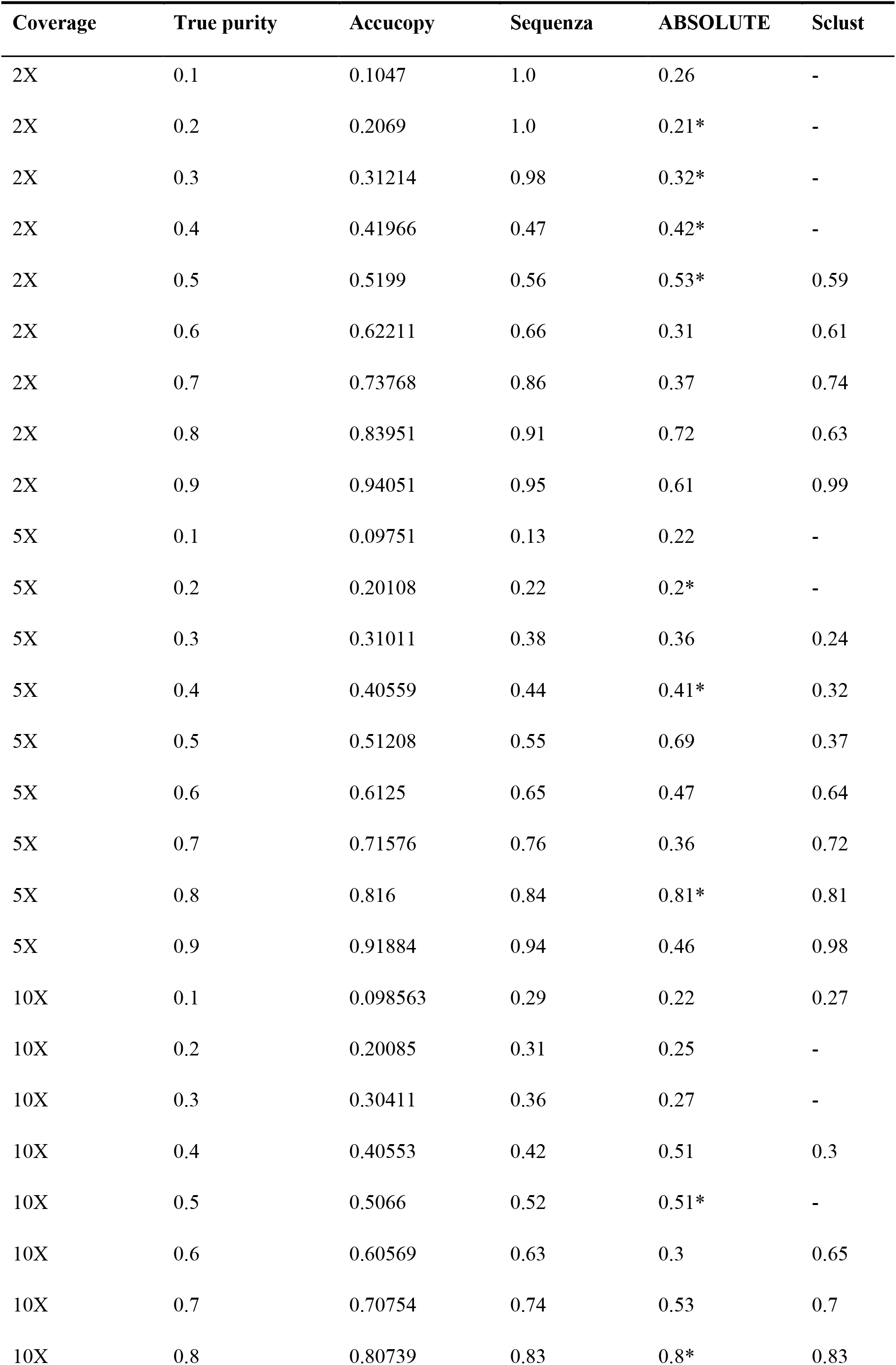
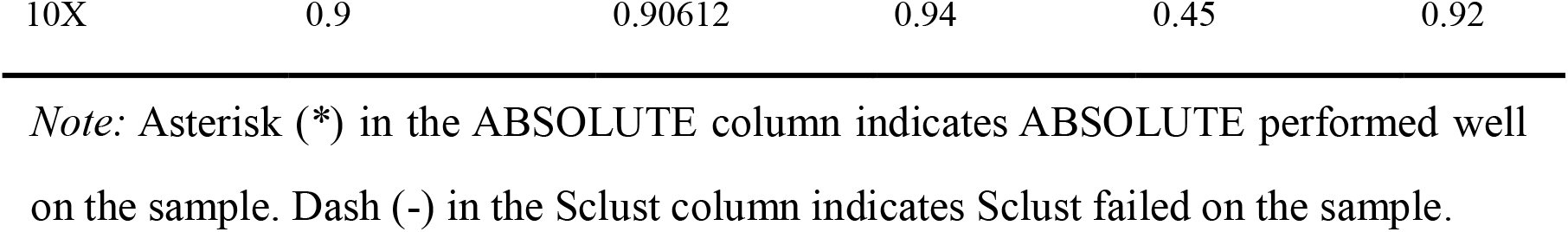
Purity estimates by all methods.

To illustrate the performance difference, we plotted the estimated TCN and ASCN of chromosome 1 by Accucopy and Sclust, with the true purity being 0.4 (low) or 0.8 (high) and coverage being 2X (low) and 10X (high) (Supplementary Fig. 1). In the 10X-coverage high-purity sample, the output by Accucopy and Sclust are very close to the truth (Supplementary Fig. 1B and 1C). If the purity decreases 0.4, the TCN and ASCN estimates of Accucopy are still very close to the truth while Sclust underestimates TCN and ASCN (Supplementary Fig. 1D and 1E). If the sequencing coverage decreases to 2X, Accucopy can still infer the true TCN in both high and low purity settings but its ASCN inference deteriorates in the low-purity low-coverage sample (Supplementary Fig. 1F and 1H). Sclust overestimated both TCN and ASCN in the 2X high-purity setting (Supplementary Fig. 1G) and failed completely in the 2X low-purity setting (Supplementary Fig 1I). This detailed comparison confirmed conclusions drawn from the summary evaluation plot (Fig. 1). The main strength of Accucopy, compared to Sclust and other methods, is that it can perform well in low-coverage and/or low-purity settings while others are unstable.

We further evaluated Accucopy on a simulated two-subclone (3:2 mixing ratio) tumor sample. The two subclones differ in six subclonal regions, colored in green, Fig. 3C. The results (Figure 3A and 3B) corroborated the single-clone simulation results. The TCN and MACN estimates by Accucopy for a purity=0.5 sample are plotted in Figure 3C and 3D. One subclonal region (chr12, copy-number=4) whose true TCN is 4=0.6*2+0.4*7, derived by averaging across the two subclones (the true copy number of this segment is 2 in subclone one and 7 in subclone two), was inferred to be clonal because Accucopy regards regions with integer copy number estimates as clonal. In reality, this is highly unlikely to happen. We designed this region to illustrate that Accucopy can recover the averaged true TCN even for subclonal regions. Accucopy only estimates MACN for clonal regions because of the infinite combinations between the number of subclones and their respective copy number states in the subclonal regions, for which Accucopy is incapable of estimating.

**Fig. 2.**
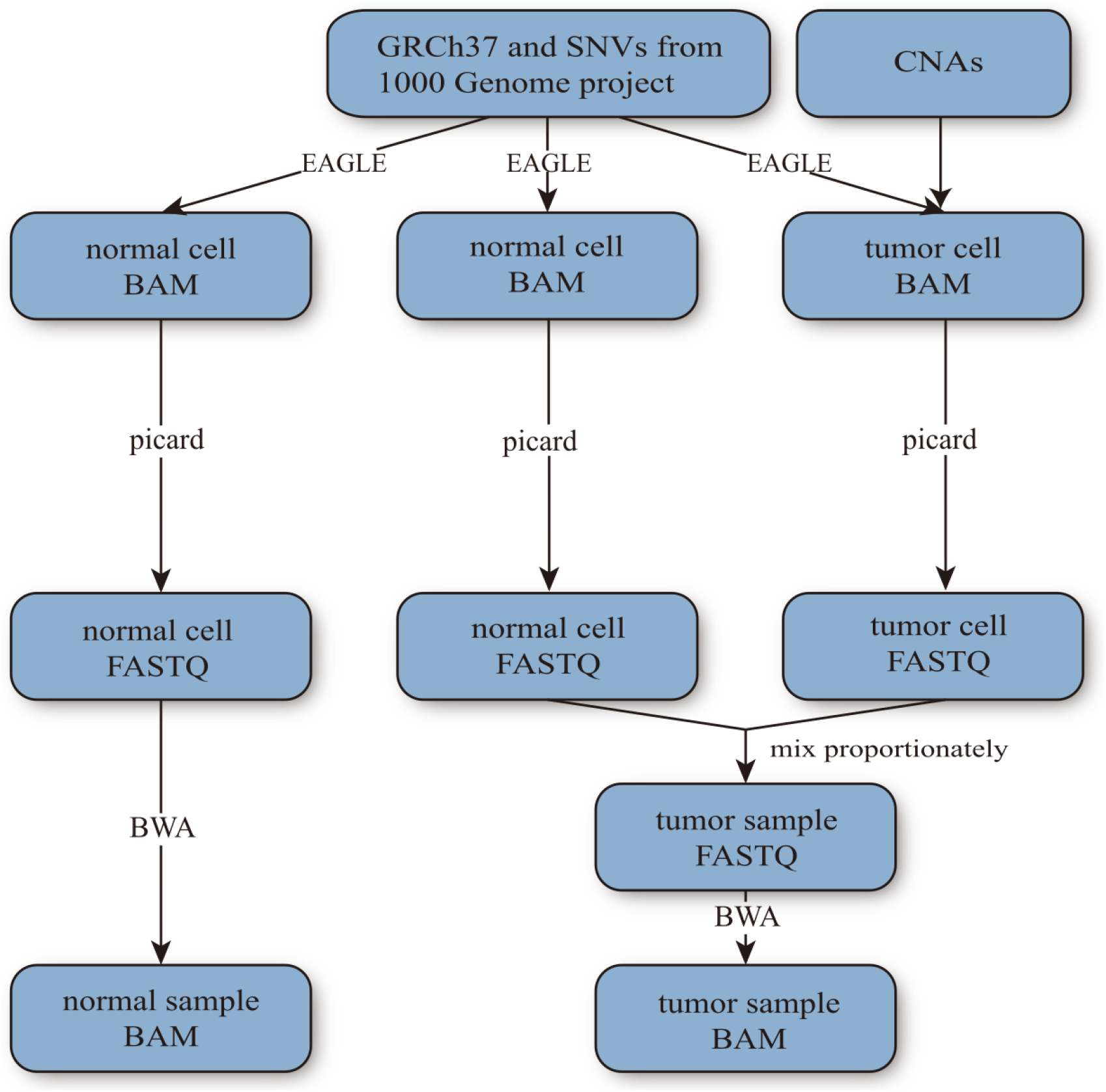
Simulation pipeline. Given the reference genome GRCh37, Single-Nucleotide-Variants (SNVs) from the 1000 Genomes project, and a set of randomly-designed CNAs to be added to the tumor cell genome, our pipeline first calls EAGLE to generate three different bam files containing simulated reads for the normal cell (the leftmost column) in the normal sample, the normal cell in the tumor sample, and the tumor cell in the tumor sample respectively. Besides random sequencing errors, EAGLE introduces point-mutations to simulated reads at the given SNV loci. Excluding the sequencing errors, the locations of the SNVs in the three different bam files are identical. The tumor cell may lose one or both SNV alleles in copy-loss regions. To simulate a tumor sample that contains subclones, we design a different set of CNAs that share some CNAs with the first tumor clone and generates another bam file for the second (or third and so forth) tumor subclone. The sequencing reads of the normal cell and tumor cell(s) that belong to the same tumor sample are mixed proportionately to generate the final bam file for one tumor sample.

**Fig. 3.**
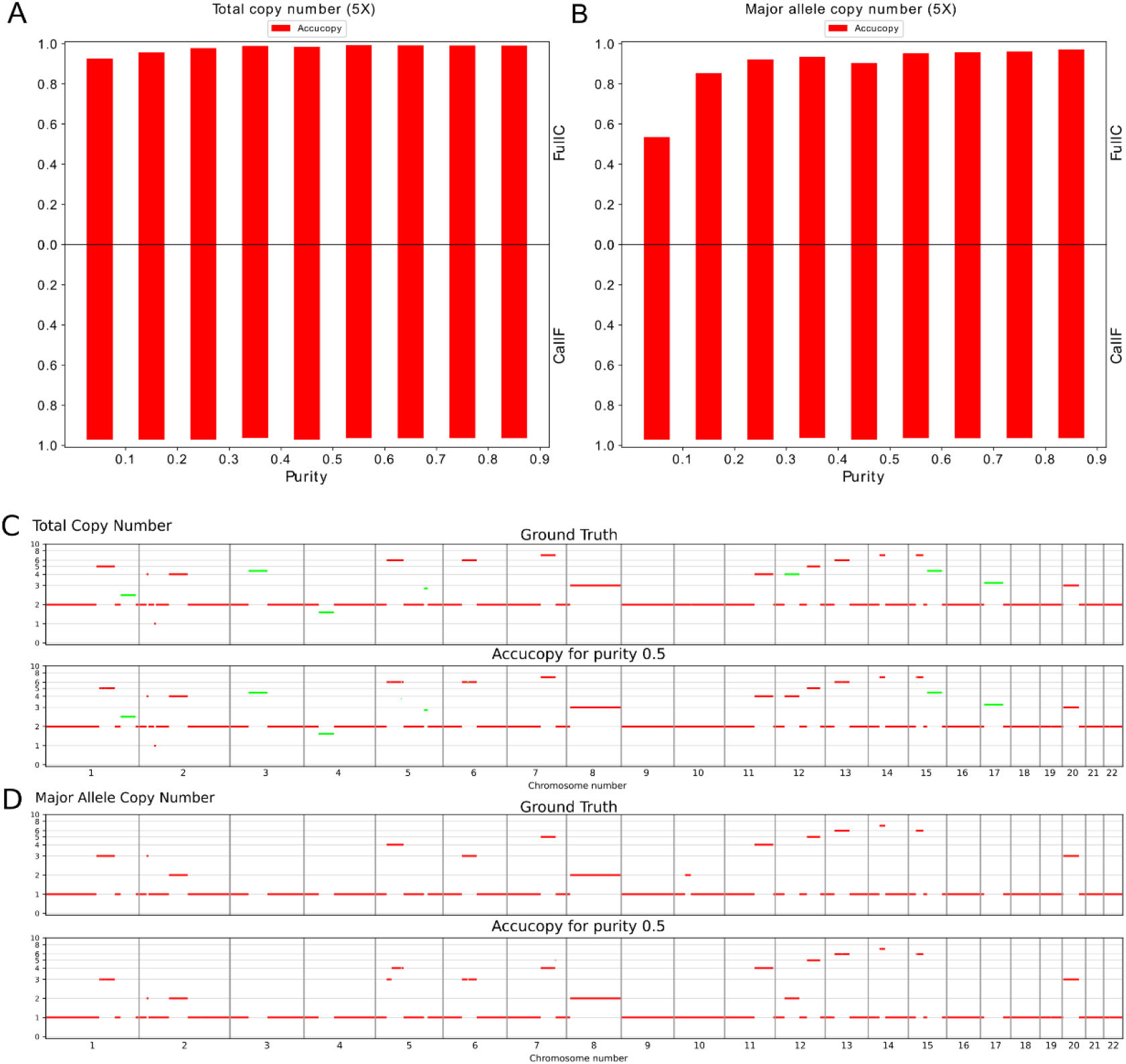
Evaluation of Accucopy on a simulated two-subclone (3:2 mixing ratio) tumor sample. The two subclones differ in six subclonal regions, colored in green in panel C. **A)** The TCN FullC results (top part of A) show the Accucopy TCN calls are at least 90% concordant with the ground truth. The TCN CallF results (bottom part of A) indicate Accucopy assigns copy numbers to close to 100% of the genome. **B)** The MACN FullC results indicate the Accucopy MACN calls are at least 85% concordant with the ground truth except when the tumor purity is below 0.1. The MACN results are very similar to those of the single-clone 5X setting. **C**) The top panel is the TCN ground truth, with seven subclonal regions colored in green. The bottom panel are the TCN estimates by Accucopy for a purity=0.5 sample. **D**) The top panel is the MACN ground truth. The bottom panel are the MACN estimates by Accucopy for the same tumor sample. Accucopy only estimates MACN for clonal regions because of the infinite combinations between the number of subclones and their respective copy number states in the subclonal regions, for which Accucopy is incapable of estimating.

The simulation analysis suggests: **a**) Accucopy can accurately estimate TCN in a wide range of purity (0.1-0.9) and coverage (2X and above) settings. **b**) Accucopy can robustly infer ASCN as long as the purity is above 0.1 in moderate or high coverage (>=5X) settings; c) In low-coverage (2X) settings, the ASCN inference by Accucopy requires the purity to be above 0.5, which suggests that a minimal 1X tumor content (=total-coverage*purity), i.e. 10X*0.1, 5X*0.2, 2X*0.5, in a sequenced sample is required for an accurate ASCN inference by Accucopy.

### Evaluation of Accucopy on the HCC1187 dataset

The prior simulation study has shown the solid performance of Accucopy in low (5X) and medium (10X) coverage settings. In this section, we run Accucopy on the HCC1187 dataset to validate Accucopy on a real sequencing dataset. We know the true purity of the eight impure HCC1187 tumor samples because we designed the mixing of HCC1187 and its corresponding normal cells. The Accucopy performance in TCN inference is on par with that of the simulation study (Fig. 4A, 4C). The MACN inference is better than that of the simulation study under similar conditions (Fig. 4B), because all the true MACNs of HCC1187 are effectively LOHs (Loss-Of-Heterozygosity). The non-LOHs of HCC1187 have unknown MACN state and are excluded in comparison. The statistical power to infer MACNs is higher for LOHs than non-LOHs because the difference between the major and the minor allele copy number is bigger for LOHs. Note that Accucopy makes MACN estimates for regions where the SKY MACN calls are missing because SKY can only call MACN for LOH regions, Figure 4D. This exercise indicates that for real-sequencing samples, Accucopy can achieve solid performance, comparable to its performance on the simulated data.

**Fig. 4.**
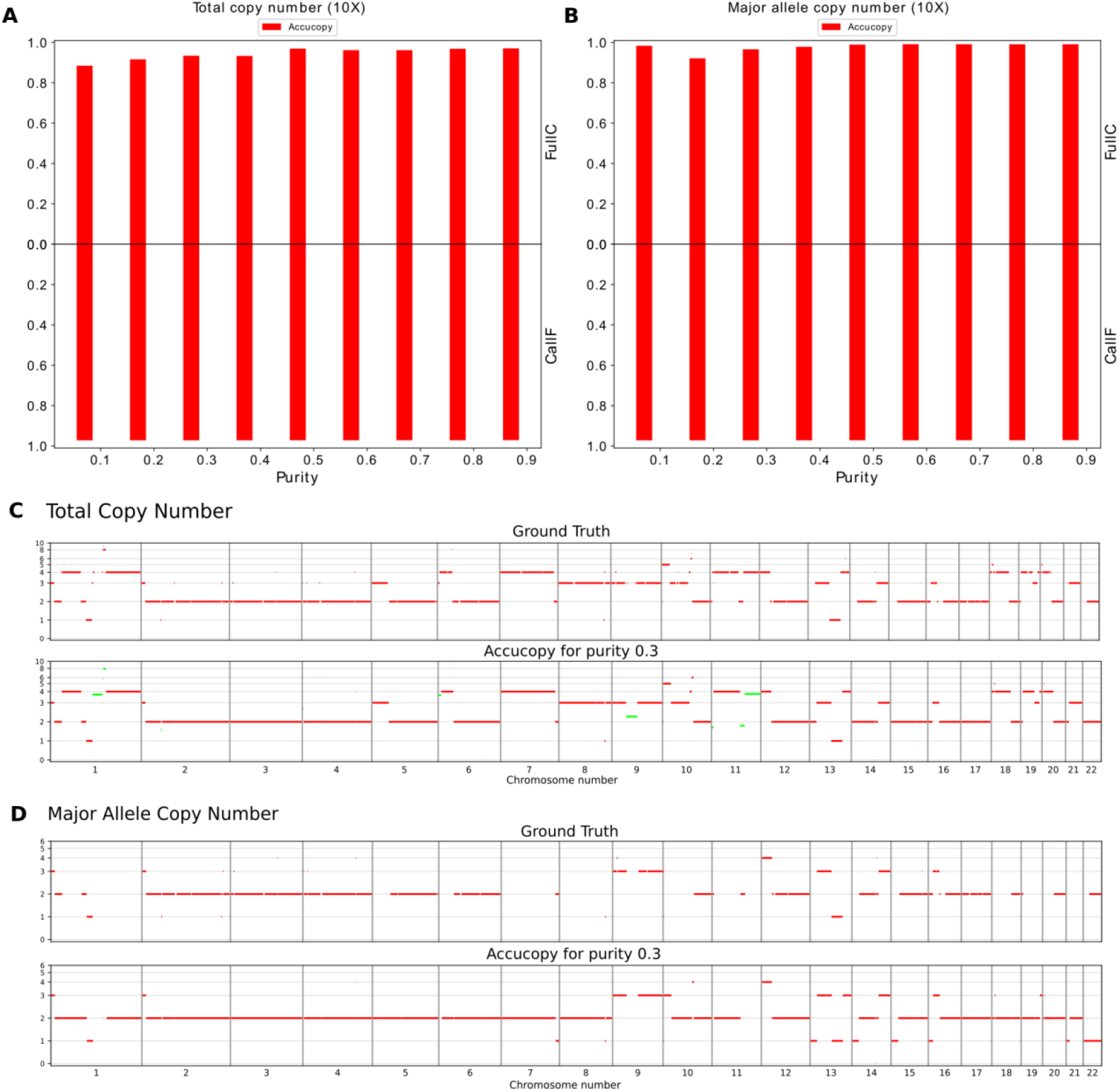
Accucopy performance on the HCC1187 dataset. The sequencing coverage for all samples is 10X. The tumor purity varies from 0.1 to 0.9. **A.** The TCN FullC results show the Accucopy TCN calls are at least 90% concordant with the ground truths. The TCN CallF results indicate Accucopy assigns copy numbers to close to 100% of the genome. **B.** The MACN FullC results indicate the Accucopy MACN calls are close to 95% concordant with the ground truth. The MACN inference is better than that of the simulation study under similar conditions because all the true MACNs of HCC1187 are effectively LOHs (Loss-Of-Heterozygosity). The non-LOHs of HCC1187 have unknown MACN state and are excluded in comparison. The statistical power to infer MACNs is higher for LOHs than non-LOHs because the difference between the major and the minor allele copy number is bigger for LOHs. **C**) The top panel is the TCN ground truth based on the spectral karyotyping (SKY) result [25]. The bottom panel are the TCN estimates by Accucopy for a purity=0.3 sample. The green segments are considered subclonal and assigned with non-integer (i.e. 3.8) copy numbers. **D**) The top panel is the MACN ground truth. The bottom panel are the MACN estimates by Accucopy for the same tumor sample. The Accucopy MACN estimates are only for clonal regions. Note that Accucopy makes MACN estimates for regions where the SKY MACN calls are missing because SKY can only call MACN for LOH regions.

### Inferring SCNAs for TCGA samples

We ran Accucopy and Sclust on 166 pairs of TCGA tumor-normal samples that have corresponding TCN profiles in the TCGA database. Accucopy succeeded for 110 samples. Accucopy failed on 56 samples due to noisy TRE data, which is caused by high level of intra-tumor heterogeneity and/or genomic alterations. Sclust succeeded for 57 samples. We compared the TCN output by either method against the corresponding TCGA TCN profiles.

The Accucopy FullC metric is strongly correlated with the tumor purity (Fig. 5A), and is independent of CallF (Fig. 5C). The average Accucopy CallF is about 95%, regardless of the tumor purity (Fig. 5B), which indicates Accucopy predicts TCNs for almost the entire genome of all analyzed samples. The Sclust FullC is also correlated with the tumor purity, but only among samples with purity above 0.5 and coverage above 10X (Fig. 6A). These samples tend to have high CallF (Fig. 6B and 6C). The decline of FullC with the decreasing tumor purity observed in both Accucopy and Sclust are quite interesting. The prior simulation and HCC1187 studies indicate that Accucopy performs well in predicting TCNs for samples with coverage 2-10X and purity 0.1-0.9 and Sclust performs well in purity>0.5 and coverage>=10X samples.

**Fig. 5.**
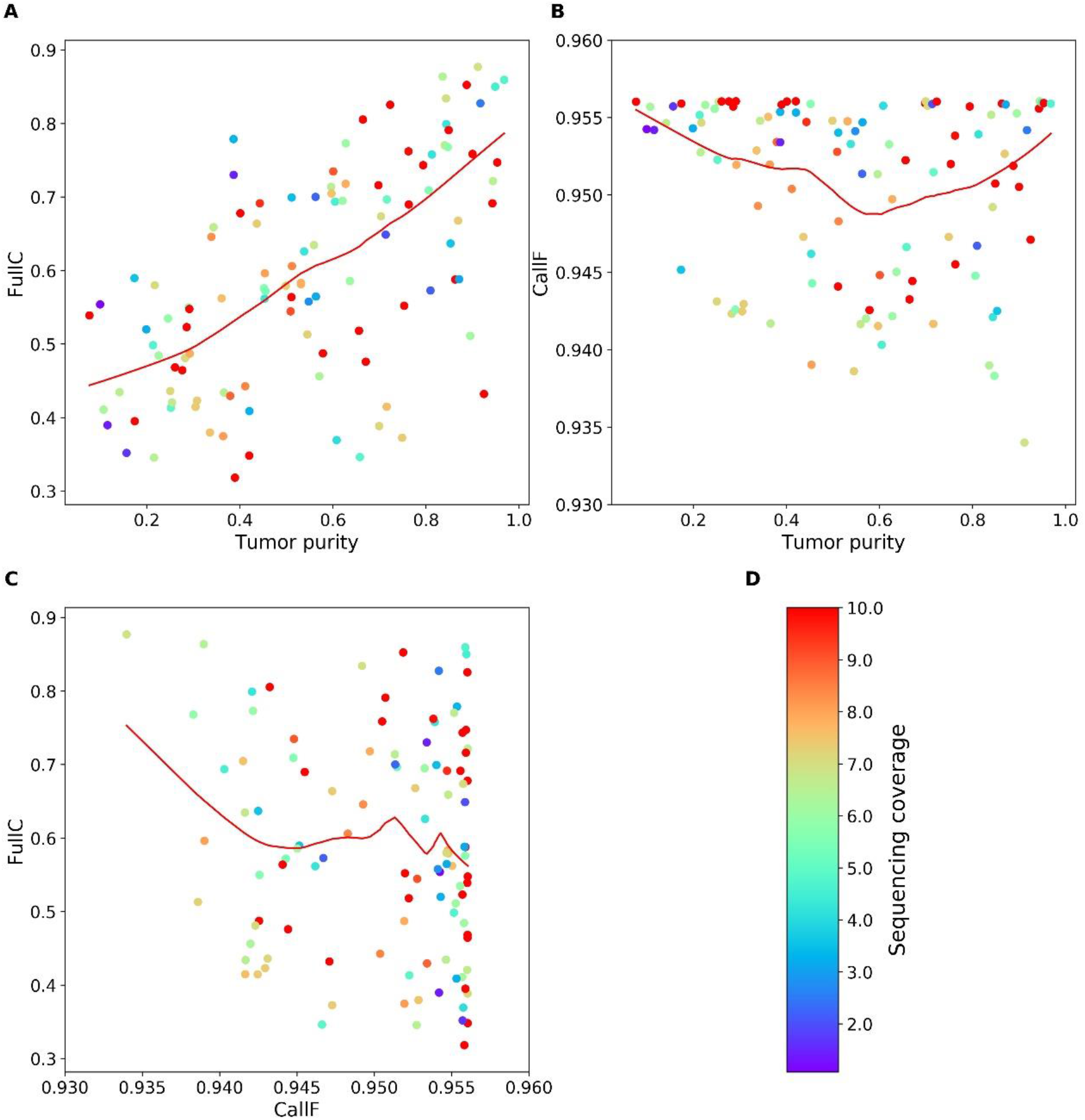
The performance of Accucopy in predicting TCNs for TCGA samples. CallF and FullC were calculated to assess the performance of Accucopy. Each dot represents one TCGA sample and is colored according to its sequencing coverage. Each scatterplot is fitted with a redline by loess smoothing. **A.** FullC is between 0.3 and 0.9, strongly dependent on the tumor purity level. **B**. CallF is between 0.93 and 0.96, independent of the tumor purity level, indicating Accucopy predicted copy numbers for almost the entire genome for all analyzed TCGA samples. **C.** FullC is independent of CallF. **D.** The colorbar maps the sequencing coverage of each sample to the color of each dot. The sequencing coverage is set to 10 for samples with sequencing coverage above 10.

**Fig. 6.**
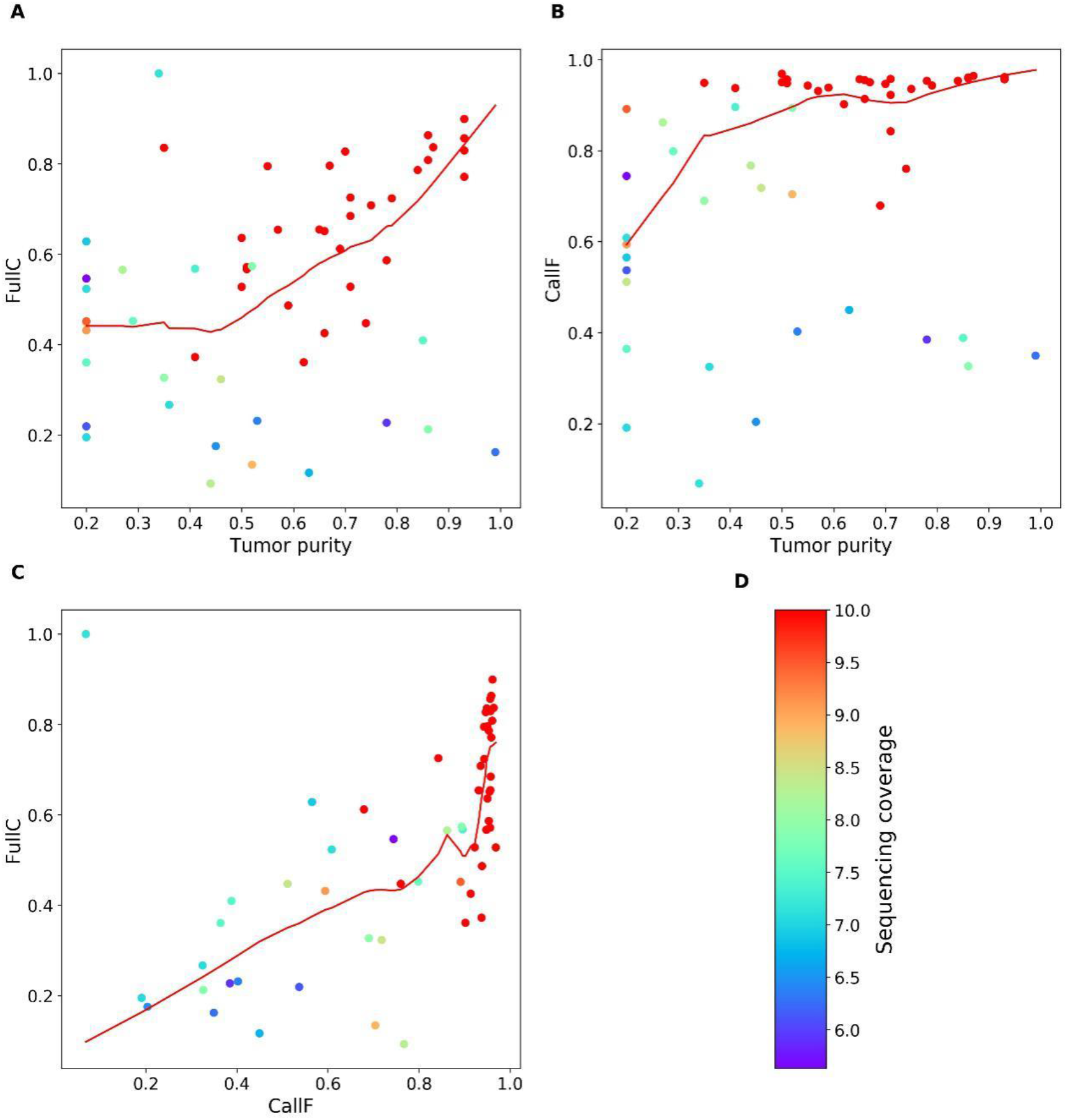
The performance of Sclust in predicting TCNs for TCGA samples. CallF and FullC were calculated to assess the performance of Sclust. Each dot represents one sample and is colored according to its sequencing coverage. Each scatterplot is fitted with a redline by loess smoothing. **A.** FullC shows a dependency on both the sequencing coverage and the tumor purity. **B.** CallF is high (~0.9) only for samples with sequencing coverage near or above 10. For samples with lower coverage, Sclust may fail to predict copy numbers for significant portions of their genomes. **C.** FullC is highly correlated with CallF. This suggests the more regions that Sclust fails to predict copy numbers, the less concordant its predicted copy numbers are with the TCGA calls. **D.** The colorbar maps the sequencing coverage of each sample to the color of each dot. The sequencing coverage is set to 10 for samples with sequencing coverage above 10. The colorbar scale starts from around 5 because Sclust failed on samples with sequencing coverage below 5.

We carefully compared the Accucopy TCN prediction for samples in the high-FullC-high-purity top-right part of Fig. 5A vs samples in the low-FullC-low-purity lower-left part of Fig. 5A and found that the decline of FullC with the decreasing tumor purity is primarily caused by the diminishing statistical power of the TCGA pipeline as the tumor purity declines (Fig. 7). The TCGA CNA pipeline (Birdsuite + CBS) assumes a tumor sample consisting of 100% tumor cells. Thus the copy number of a genomic segment predicted by the TCGA pipeline is a weighted average of its respective copy numbers in the tumor and normal cells. As the purity of a tumor sample declines, the increasing fraction of normal cells, whose genomic copy number is two, will move the predicted average copy number closer to two. Accucopy and Sclust explicitly model the tumor purity and do not suffer from this issue. This is shown in detailed TCGA vs. Accucopy comparisons (Fig. 7). In both samples, the segmentations of the genome by the TCGA pipeline and Accucopy are highly similar. In addition, the copy number qualitative predictions (duplication or deletion) for individual segments are highly similar too. Were it not for FullC to consider copy number differences, both samples would have shown near perfect concordance between the TCGA profile and the Accucopy output. In the low-purity sample (Fig. 7A), the copy number quantitative predictions of abnormal segments are closer to two and are numerically less concordant with those by Accucopy, manifested by a lower FullC than the high-purity sample (Fig. 7B).

**Fig. 7.**
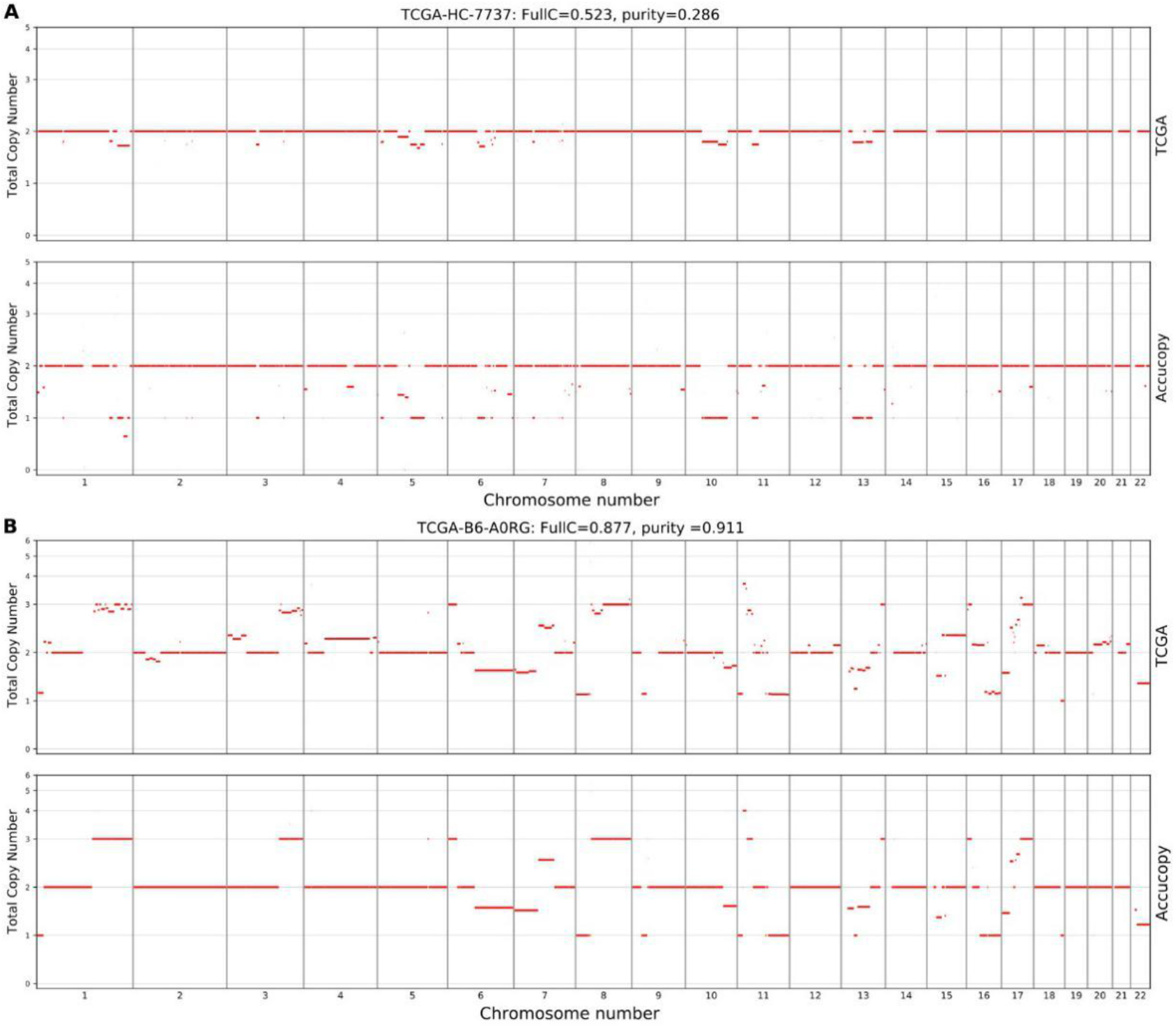
The loss of power of the TCGA CNA pipeline in low-purity samples. The copy number of a genomic segment predicted by the TCGA pipeline, which does not model the tumor purity, is a weighted average of its respective copy numbers in the tumor and normal cells. As the purity of a tumor sample declines, the increasing fraction of normal cells will move the predicted average copy number closer to two. Accucopy treats the copy numbers of the tumor and normal cells within one tumor sample as two separate parameters in its model. Panel (**A**) exhibits the copy number profile of a low-purity sample (purity=0.286) predicted by the TCGA pipeline, the upper panel, versus that predicted by Accucopy, the lower panel. The FullC between the two profiles is 0.523. Panel (**B**) is a similar plot to panel A, for a high-purity sample (purity=0.911). The FullC between the TCGA-pipeline and Accucopy predicted CNA profiles is 0.877. In both samples, the segmentations of the genome by the TCGA pipeline and Accucopy are highly similar. In addition, the copy number qualitative predictions (duplication or deletion) for individual segments are also highly similar. However, in the high-purity sample (B), the copy number quantitative predictions of abnormal segments are further away from two and are numerically more concordant with those by Accucopy, manifested by a higher FullC than the low-purity sample (A).

It is also clear from Fig. 6 that Sclust works in more limited conditions than Accucopy. Sclust requires the tumor purity above 0.5 and the sequencing coverage above or near 10X. For samples with sequencing coverage below 10X, Sclust may predict copy numbers for only a fraction of the genome (Fig. 6B). For samples with coverage lower than 5X, (Fig. 6D), Sclust failed completely. This is consistent with the simulation finding that Sclust has lower power in detecting CNAs from the low-purity (<0.5) and/or low-coverage (<=5X) samples.

The TCGA study indicates Accucopy is capable of identifying copy number alterations in complex real-world samples, some of which may have very low sequencing coverage and are of low tumor purity.

### Implementation and performance

Accucopy is implemented in vanilla C++ and Rust and is released for Ubuntu 18.04 in a docker. In theory, it can be built for Windows or MacOS but we have not tested it. Average runtime of Accucopy is about one hour for a 5X tumor/normal matched pair; about four hours for a 30X tumor/normal matched pair on a single core of Intel(R) Xeon(R) CPU E5-2670 v3 @ 2.30GHz; with the peak RAM consumption under 4GB.

We provided all methods with the same input bam files and ran all programs with default parameters under the same computational environment as stated above.

## Discussion

Through extensive simulated and real-sequencing data analyses, we have demonstrated that Accucopy is a fast, accurate, and fully automated method that infers TCN and ASCN of somatic CNAs from tumor-normal high-throughput sequencing data. The strength of Accucopy, relative to other methods, lies particularly in its performance in low-coverage and low-purity samples. This makes Accucopy an excellent choice in first-round low-coverage screening type of analysis. It can offer crucial insight regarding the tumor purity, ploidy, TCNs, and ASCNs before an expensive in-depth high-coverage analysis is started.

One under-appreciated factor contributing to the excellent performance of Accucopy is the large amount of simulation and real-sequencing samples with known truth (or near truth for TCGA samples). This trove of data leads us to adjust many aspects of the Accucopy model during development. Here are a few notable adjustments. A coverage smoothing step greatly reduced the random noise in sequencing coverage. Adoption of Strelka2 [22] dramatically reduced the number of false positives in calling heterozygous SNPs, compared to other variant callers we tried. Extensive in-depth analyses uncovered that the expectation of Log ratio of Allelic Ratios (LAR) needed to be adjusted due to the exclusion of zero-allele-coverage SNPs, which improved the Accucopy performance in the ASCN inference by an order of magnitude. These adjustments look trivial but cumulatively are very effective in improving the overall performance of Accucopy.

The requirement of a periodic TRE pattern arising from varying copy numbers means that Accucopy is not suitable for tumor samples with little or no copy number alterations. An excessive amount (i.e. >3%) of point mutations in a tumor, relative to its matching normal, causing many wrong alignments, will also render Accucopy unable to confidently discover a period from the TRE pattern because the input coverage information is no longer valid. Another case that could weaken Accucopy is the presence of copy number variations (CNVs) in healthy normal individuals. At these genomic regions, the TCN and ASCN predictions by Accucopy will be inaccurate as Accucopy assumes the entire genome of a normal sample to be of copy number two. For intermediate-purity tumor samples, Accucopy can discover events between 1-10Mb, as demonstrated in Figure 3C and Figure 4C. However for low-coverage and/or low-purity samples, we suspect Accucopy will perform poorly on focal events (far less than 1Mb) because these regions will provide little data for accurate statistical inference. In this kind of scenario, the Accucopy result can form the basis for the next-round decision-making. For example, if a clinical tumor sample is of very low purity (<0.1) based on a first-round shallow sequencing, scientists may choose another higher-purity tumor sample for high-coverage sequencing because the latter contains more information.

Whole-genome duplication (WGD), quite prevalent in cancer genomes, if happening cleanly, i.e. copy 2 becoming copy 3, offers no discernable periodic TRE pattern for Accucopy to analyze and thus Accucopy will be incapable of doing any inference as there will be only one peak of genomic coverage. However, in real tumor genomes, WGDs are rarely that clean and usually involve many gains and losses and the copy number differentiation in different genomic regions provides opportunity for Accucopy to make inference. The HCC1187 genome, Figure 4C, is an example in which a WGD event has gone awry causing many gains and losses. As shown in Figure 4, Accucopy is quite capable to making inference from these kinds of WGD-ed samples.

For regions where different tumor subclones harbor different SCNAs, producing averaged TCNs and MACNs is less than satisfactory from the standpoint of tumor clonal evolution. We are currently developing the next iteration of Accucopy to address this issue.

## Conclusions

Through extensive simulated and real-sequencing data analyses, we have demonstrated that Accucopy is a fast, accurate, and fully automated method that infers TCN and ASCN of somatic CNAs from tumor-normal high-throughput sequencing data. The strength of Accucopy is particularly in its performance in low-coverage low-purity samples. This makes Accucopy an excellent choice in first-round low-coverage screening type of analysis. It can offer crucial insight regarding the tumor purity, ploidy, TCNs, and ASCNs before an expensive in-depth high-coverage analysis.

## Methods

### Simulated tumor and matching normal sequencing data

We generated *in silico* tumor and matching-normal WGS data using an EAGLE-based workflow at three coverage settings: 2X, 5X, and 10X. [EAGLE](https://github.com/sequencing/EAGLE) is a software developed by Illumina to mimic their own high-throughput DNA sequencers in terms of sequencing biases and errors. We introduced twenty-one somatic copy number alterations (SCNAs), with length ranging from 5MB to 135MB and copy number from 0 to 8, affecting about 28% of the genome, to each simulated tumor genome. The entire genome of its matching normal sample is of copy number two. Over one million heterozygous single-nucleotide loci (HGSNVs) were introduced to each normal and its matching tumor sample. For each coverage setting, we first generated a pure tumor sample (purity=1.0) and its matching normal sample. We then generated nine different impure tumor samples (purity from 0.1 to 0.9) by mixing the pure tumor sample sequencing with its matching normal data proportionately. The mixing proportion determines the tumor sample’s true purity. The simulation pipeline for one pair of tumor-normal samples is visualized in Figure 2.

### HCC1187 cancer cell line dataset

The genome-wide CNA profile of HCC1187 has been widely studied via the spectral karyotyping (SKY) tool, which is one of the most accurate tools for characterizing and visualizing genome wide changes in ploidy [23, 24]. We used the SKY result from [25] as the ground truth for CNA comparison. SKY does not reveal a genome-wide ASCN profile and only identifies the LOH (Loss-of-Heterozygosity) regions. For these LOH regions, about 60% of the HCC1187 genome, we inferred the ASCN based on their LOH and CNA states. The whole genome sequencing data of pure HCC1187 cancer cells and its matched normal HCC1187BL cell lines was downloaded from Illumina BaseSpace. The sequencing coverage for HCC1187 and HCC1187BL is 104X and 54X respectively. Based on the pair of pure-tumor and normal real sequencing data, we generated eight impure tumor samples with purity from 0.1 to 0.9 (exclude 0.5) by proportionately mixing the HCC1187 reads with its matching normal reads.

### TCGA samples

One hundred sixty-six random pairs of TCGA tumor-normal samples, were downloaded from TCGA to provide a more comprehensive evaluation of Accucopy on real-world samples. The cancer types include breast invasive carcinoma (BRCA), Colon adenocarcinoma (COAD), Glioblastoma multiforme (GBM), Head and Neck squamous cell carcinoma (HNSC), and Prostate adenocarcinoma (PRAD). The TCGA database also contains CNA profiles that are derived from Affymetrix SNP 6.0 array data. The TCGA CNA calling pipeline is built onto the existing TCGA level 2 data generated by Birdsuite [26] and uses the DNAcopy R-package to perform a circular binary segmentation (CBS) analysis [27], which translates noisy intensity measurements into chromosomal regions of equal copy number. These TCGA database CNA profiles are only for TCN (Total Copy Number). During comparative analysis, we compared Accucopy TCN estimates with these TCGA database TCN estimates.

### Evaluation metrics

Define T and P as the truth and the predicted sets of copy-number segments of a sample respectively.

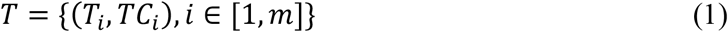

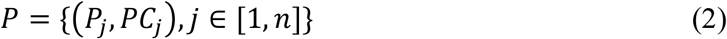

In equations above, m and n are the number of the segments in the truth and predicted sets respectively, *T_i_* or *P_j_* is the coordinate interval of a segment in the form of (chromosome, start, stop), and *TC_i_* or *PC_j_* is the copy number (float type) of this segment. Segments with no copy-number assigned by a method are excluded from *T* and *P* because any normal or abnormal assumption regarding their copy number state is hard to justify.

To evaluate the performance of a method, we defined two metrics. The first metric, CallF, is the fraction of the genome whose copy number state is assigned by a method.

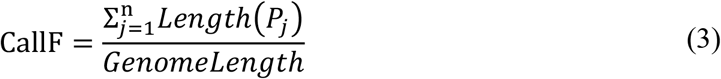

The second metric, FullC, Eq. 4, is a correlation-like metric that measures how the predicted CNAs are concordant with the truth set, considering both the segment coordinates and the copy-number difference. For one predicted segment, *P_j_*, FullC finds all truth segments that intersects with it. No threshold is involved in finding intersections, thus any overlap is an intersection. One predicted segment could have multiple intersections with the truth set. The denominator is the sum of the length of all intersections, which is usually less than the genome length because the normal (copy-number=2) segments are excluded (details below). The numerator is the sum of the product between the length of an intersection, *Length*(*T_i_* ∩ *P_j_*), and the copy-number-difference distance metric, *e^−|TC_i_−PC_j_|^*. The latter is an application of the exponential function to the absolute copy-number difference, thus converting the difference to a distance metric with its value in the range of 0 and 1. Segments that are normal (copy-number=2) in both *T* and *P* are excluded. If normal segments are part of the comparison, in cases where copy number 2 is sometimes the most ubiquitous state of a cancer genome, a simple method calling the entire genome as copy number 2 will show good results.

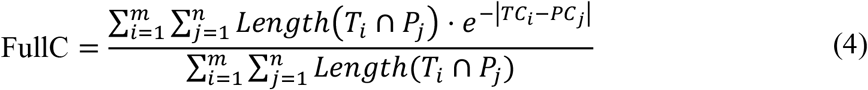

The value of FullC is between 0 and 1, with 0 being completely discordant and 1 being completely concordant and can be applied to both TCN and ASCN performance evaluation. In the ASCN case, the FullC for the Major Allele Copy Number (MACN) is equal to the FullC for the minor allele copy number and we only show the MACN FullC. Notably, FullC treats large copy number aberrations and focal amplifications and losses equally. The evaluation metric used by early publications that ignores the mismatch of copy numbers and only considers the concordance of the coordinates of non-normal segments, can be misleading. For example, a duplication could be considered a match to a deletion. FullC remedies this issue by considering the copy number difference.

For the simulation data and the HCC1187 dataset, the truth set is known. For the TCGA samples, we use the TCN profiles in the TCGA database as a proxy for the truth set. They are derived from the Affymetrix SNP 6.0 array, not strictly a truth set, but helpful in our comparison analyses.

### Summary for the Accucopy model

The Accucopy model is a probabilistic model that infers the TCN and ASCN from two types of input: the sequencing coverage information summarized by Tumor Read Enrichment (TRE) and the allele-specific coverage information summarized by Log ratio of Allelic-coverage Ratios (LAR) at HGSNVs. Definitions of TRE and LAR are in Additional File 1. TREs are samples from a multi-component Gaussian mixture model with the tumor purity, the tumor ploidy, and the total copy number of each genomic region as parameters. LARs are samples from a two-component Gaussian mixture model with the allele-specific copy numbers of genomic regions as additional parameters. The entire genome is segmented by an enhanced version of the public GADA [28] (unpublished). We developed an autocorrelation-guided EM algorithm to find optimal parameters. Bayesian Information Criterion is adopted to avoid model overfitting. More details of the Accucopy model are in the Additional File 1.

## Availability and Requirements

Project name: Accucopy

Project home page: https://github.com/polyactis/Accucopy Operating system(s): Linux

Programming language: C++/Rust/Python

License: SIMM Institute License, free for non-commercial use.

Any restrictions to use by non-academics: license needed.

## Abbreviations

CNAs: Copy number alterations
TCN: total copy number
ASCN: allele-specific copy numbers
SCNAs: somatic copy number alterations
HGSNVs: heterozygous single-nucleotide loci
SKY: spectral karyotyping
LOH: Loss-of-Heterozygosity
BRCA: breast invasive carcinoma
COAD: Colon adenocarcinoma
GBM: Glioblastoma multiforme
HNSC: Head and Neck squamous cell carcinoma
PRAD: Prostate adenocarcinoma
CBS: circular binary segmentation
MACN: Major Allele Copy Number
TRE: Tumor Read Enrichment
LAR: Log ratio of Allelic-coverage Ratios
CNVs: copy number variations

## Declarations

### Ethics approval and consent to participate

This publication only uses public data only. Therefore, we do not need ethics approval or consent to participate.

### Consent for publication

Not applicable.

### Availability of data and materials

The datasets analyzed during the current study are available in the TCGA. Barcode and UUID of each TCGA sample file can be found in Additional File 3.

### Authors’ contributions

YSH conceived the idea and supervised the study. XF and YSH designed and implemented the method. XF and GL performed the simulated data analysis. XF performed the other data analyses. XF and YSH wrote the manuscript. All authors read and approved the final manuscript.

## Acknowledgements

We wish to acknowledge the guidance provided by the authors of Sclust on how to configure their software.

## Supplementary information

**Additional File 1:** Method details of Accucopy.

**Additional File 2: Supplementary Figure 1.** Copy number profile of chr1 on partial simulation data. X-axis is the chromosomal position and Y-axis is the copy number. Each figure has two panels. The top is the profile of absolute copy number and the bottom is the profile of major allele copy number. **A.** The truth profile of chr1. **B-I.** The copy number profile given by different methods on different samples. The title of each figure has three keys split by space, which indicate the method name, sample coverage and sample purity respectively.

**Additional File 3:** Barcode and UUID of each TCGA sample.

